# Local, Quantitative Morphometry of Fibroproliferative Lung Injury using Laminin

**DOI:** 10.1101/2023.06.15.545119

**Authors:** Brendan P. Cox, Riley T. Hannan, Noora Batrash, Jeffrey M. Sturek

**Affiliations:** Department of Medicine, Division of Pulmonary and Critical Care Medicine, University of Virginia School of Medicine; University of Virginia School of Medicine

## Abstract

Investigations into the mechanisms of injury and repair in pulmonary fibrosis require consideration of the spatial heterogeneity inherent in the disease. Most scoring of fibrotic remodeling in preclinical animal models rely on the modified Ashcroft score, which is a semi-quantitative scoring rubric of macroscopic resolution. The obvious limitations inherent in manual pathohistological grading have generated an unmet need for unbiased, repeatable scoring of fibroproliferative burden in tissue. Using computer vision approaches on immunofluorescent imaging of the extracellular matrix (ECM) component laminin, we generate a robust and repeatable quantitative remodeling scorer (QRS). In the bleomycin lung injury model, QRS shows significant agreement with modified Ashcroft scoring with a significant Spearman coefficient r=0.768. This antibody-based approach is easily integrated into larger multiplex immunofluorescent experiments, which we demonstrate by testing the spatial apposition of tertiary lymphoid structures (TLS) to fibroproliferative tissue. The tool reported in this manuscript is available as a standalone application which is usable without programming knowledge.

## Introduction

Interstitial lung diseases (ILD) comprise a diverse group of lung pathologies with varying drivers of tissue injury ultimately leading to aberrant tissue remodeling and scar. Idiopathic pulmonary fibrosis is an ILD typified by chronic, progressive interstitial fibrosis with a characteristic histologic appearance^1^. Loss of lung function due to fibroproliferation results in mean survival time of 2-5 years after diagnosis. The cause of IPF is unknown and is thought to be a combination of genetic and environmental factors^2^.

Histopathologic analyses of IPF, presenting as usual interstitial pneumonia (UIP), are a core component of ongoing research into the mechanisms driving fibrotic disease. The pathologist’s diagnosis of UIP as described by guidelines updated in 2018 require 1) normal lung parenchyma punctuated with regions of fibrotic remodeling, 2) fibroblastic foci (myxoid, pale staining subepithelial fibroblasts), 3) accumulation of ECM/scar, and 4) honeycombing (dilated cystic spaces lined by airway epithelium located within mature scar), in addition to the absence of features suggesting other diagnoses^3,4^. These fibroblastic foci can be observed at the boundary between uninvolved tissue and advancing scar and are where tissue remodeling and scar-generating myofibroblasts proliferate^5^. Effective accounting for this spatially variable biology is therefore an important factor in the study of pathologic mechanisms^6,7^. Adequately recapitulating and studying these phenomena in preclinical animal models is an evergreen problem for those studying lung fibrosis The Ashcroft scoring system, put forward by Ashcroft et.al. in 1988^8^ and modified further by Hübner et.al. in 2008^9^, is an expansion on even earlier disease staging guidelines for asbestosis^10^, and consists of eight grades of fibrosis and a score of zero for healthy lung. The scoring is tacitly performed on lung parenchyma, with contributions from airways and larger blood vessels excluded heuristically by pathologists. Grades one through four are characterized by thickening of interstitial tissue but no loss of alveolar architecture. Grades five through eight are all characterized by distinct fibrotic masses and progress in severity with the proportion of parenchyma displaced by tissue, up to complete ablation of normal tissue at a score of 8. This scoring is traditionally performed on sections stained with Masson’s trichrome, which emphasizes collagen content, but this is not required.

For several reasons, modified Ashcroft is not conducive to multiplexed imaging approaches. It requires histochemical staining, which is generally incompatible with multiplex immunofluorescent imaging. It is traditionally scored on large fields of view, limiting the resolution of fibrotic scoring to gross evaluations. Finally, it is performed manually, introducing bias and variability. There are computational methods in active development which automate the histopathologic scoring of chemical stains, successfully removing the burden of labor and inter/intra-observer variability from the analysis, but these methods do not provide the capability to multiplex with other targets^11,12^.

Immunofluorescent approaches benefit from target specificity and ease of multiplexing, but are generally deficient in broader tissue context, as only targeted antigen are visible. Frequently, extracellular matrix proteins such as collagens, laminins, or elastins are targeted in multiplex immunofluorescent panels to provide this missing structural information^13^. A distinct structure in healthy tissue is the basement membrane, or basal lamina; a thin sheet of extracellular matrix (ECM) upon which cells are anchored and derive their polarity. To maximize gas exchange in the lung, the alveolar basement membrane is quite thin and is frequently shared between capillary endothelium and alveolar epithelium. The primary structural components of this basement membrane are laminin and collagen IV. The loss of organized basement membrane structure is thought to mark irreversible lung injury; functional tissue has been remodeled by fibroproliferative processes such that the blueprint for healthy tissue has been lost^14^. While laminin and/or collagen IV are frequently targeted for immunostaining, there is an unexploited opportunity to extract further information about tissue health from these basement membrane proteins.

The tool described in this study uses analyses of immunofluorescent signal from a polyclonal anti-laminin antibody to compute a local quantitative remodeling score (QRS). The morphological progression of the laminin network during fibroproliferation can be described as a broadening of the cross-sections of alveolar basement membrane, and loss of contrast between basement membrane and adjacent tissue. These qualitative phenomena are found concomitant with reduction in alveolar air space and expanding interstitial space in the tissue. These changes in morphology are easily visible to the human observer but require textural image analysis to be quantified, a field of image processing common in medical imaging. Herein, we validate our tool against the common histopathologic scoring rubric of modified Ashcroft in the bleomycin preclinical animal model, and demonstrate a novel spatial analysis as a use case enabled by this method.

## Methods

Detailed methods for use of the QRS app are described in the supplement.

### Preclinical Animal Model of Bleomycin Lung Injury

8-12-week old C56BL/6J mice (The Jackson Laboratory) were anesthetized in accordance with IACUC approved protocols. Clinical grade bleomycin sulfate (Pfizer/Fresenius Kabi via University of Virginia Health System Pharmacy Services) was introduced orotracheally and aspirated at a dose of one unit per kilogram in a volume of 50ul of sterile saline. To capture fibroproliferation, mice were euthanized at two weeks-post bleomycin treatment.

### Sectioning and Histology of Lungs

Mice were perfused, lungs lavaged once with saline then inflated with 1% low melting temperature agarose (Invitrogen) in 1xPBS. Lungs were fixed in 4% paraformaldehyde in 1xPBS, then cryoprotected in 30% sucrose in 1xPBS until sinking. Lungs were dissected into lobes, embedded in OCT and frozen. The left lobe was sagittally sectioned at 10 μm thickness onto Superfrost Plus slides (Fisher Scientific).

### Histochemistry, Immunostaining, and Imaging

Sections were permeabilized and blocked as described previously^15^. Anti-col1a1 (AB_2904565, Cell Signaling Technology), anti-laminin (AB_10001146, Novus), anti-CD45R/B220 (AB_2896201, eBioscience) for B cells, and dapi or NucSpot Live 488 (Biotium) were used for nuclear staining. Stained samples were mounted in SlowFade Glass (Invitrogen). Micrographs were acquired on a Leica Thunder TIRF instrument in epifluorescence (Leica DMI8), using 20x objective (Leica HC PL APO 20x/0.80 DRY). Laminin fluorescence was corrected for flatness of field using a dyed slide reference and stitched using LAS-X (LAS X 3.7.5.24914).

For sequential staining of sections, initial immunostaining and imaging was performed as described, the samples de-coverslipped, and then stained for Masson’s trichrome

### Image Analysis

For the standalone application, re-exported TIFF images were batch processed with the parameters listed in the supplement. Images were loaded into QuPath and overlaid if necessary. For modified Ashcroft scoring, ten random fields of 900 microns square were taken from each whole slide Masson’s trichrome scan, with four blinded scorers scoring all fields in a single session. For the correlation of computed features to modified Ashcroft scores, the average computed features within the same full-sized Ashcroft-scored field were compared. A full list of correlated features is available in the Supplement. For all immunofluorescence analysis and visualization, true signal was determined using unstained and secondary-only controls. The laminin channel was used with the thresholding function in QuPath to generate masks of lung tissue. TLS were manually identified and distance maps between TLS and tiles generated using the QuPath function distanceMap. Measurements were exported, with subsequent manipulations and statistical tests performed in Graphpad Prism 9.0.

## Results

### Basement membrane remodeling in bleomycin model of fibroproliferative lung disease

The orotracheal bleomycin injury model for fibroproliferative disease induces collagen deposition, myofibroblast activation and proliferation, and destruction of functional lung parenchyma^16,17^. At fourteen days post-bleomycin administration, the acute inflammatory phase is largely resolved, and instead fibroproliferation predominates, with maximal scar appreciable at 21-28 days post-injury. Here, we look at early tissue remodeling, with the goal of identifying features associated with fibroproliferation prior to the generation of mature scar.

Heterogeneity of injury phenotype is apparent within a single section at low magnification (Figure 1A) with fibroproliferative (gold box) and healthy (cyan box) tissue delineated and seen in greater detail in Figure 1B. The deposition of collagen in parenchymal space, visualized blue via Masson’s trichrome (Figure 1A, left) or with anti-collagen antibody (Figure 1A, center) indicates areas of fibroproliferation. Immunostaining for collagen alone does not distinguish between preexisting healthy collagen, as it is abundant in the cuff spaces of healthy airways (Figure 1B asterisks), and pathologically deposited in the interstitium in fibrosis. The addition of laminin immunostaining provides structural context (Figure 1A, B, right) and alveolar spaces become readily identifiable. Laminin alone is sufficient to qualitatively identify tissue remodeling in lung parenchyma: healthy networks of alveolar basement membrane exhibit a characteristic thin and highly networked structure, while the same basement membrane network becomes broad and loses contrast in regions of remodeling (Figure 1C top and bottom, respectively).

**Figure 1.**
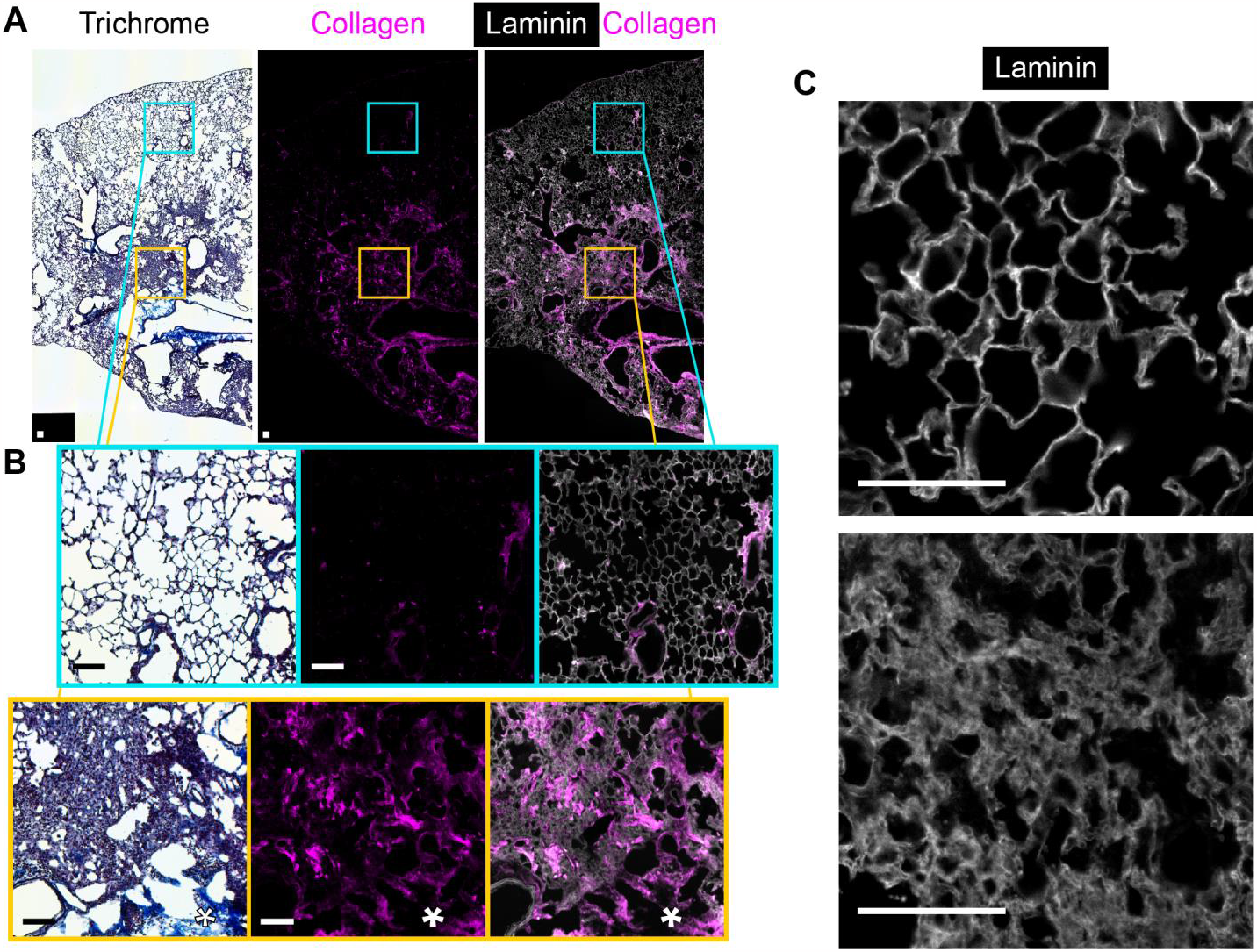
Basement Membrane Remodeling Phenotype in Bleomycin Lung Injury. **(A)** Representative micrographs of serial sagittal sections of a left mouse lung two weeks post bleomycin treatment. Histochemical staining for Masson’s trichrome **(A, left)** and immunofluorescence for collagen I (anti-Col1a1) alone **(A**,**middle)** and with laminin (anti-laminin) **(A, right). (B)** Expanded insets of micrographs, taken from healthy **(B, top)** and fibroproliferative **(B, bottom)** regions of the same section. Asterisk indicates the collagen-rich, laminin-negative cuff space surrounding large airways. **(C)** Higher resolution fluorescence micrographs of laminin from a serial section of the same lung. Field from healthy **(C, top)** and fibroproliferative (**C, bottom)** regions of lung parenchyma. Scale bars = 100 microns.

### Generation of computed laminin features and association with modified Ashcroft

Sequential staining of a single section allows for the 1:1 alignment of multiple imaging modalities performed in series. Initial processing for immunofluorescence imaging is performed as normal. After fluorescence imaging, the sample is de-coverslipped, processed for Masson’s trichrome staining, and imaged in color via brightfield microscopy (workflow schematized in Figure 2A). Each lung section has now been imaged twice: laminin immunofluorescence and color Masson’s trichrome.

**Figure 2.**
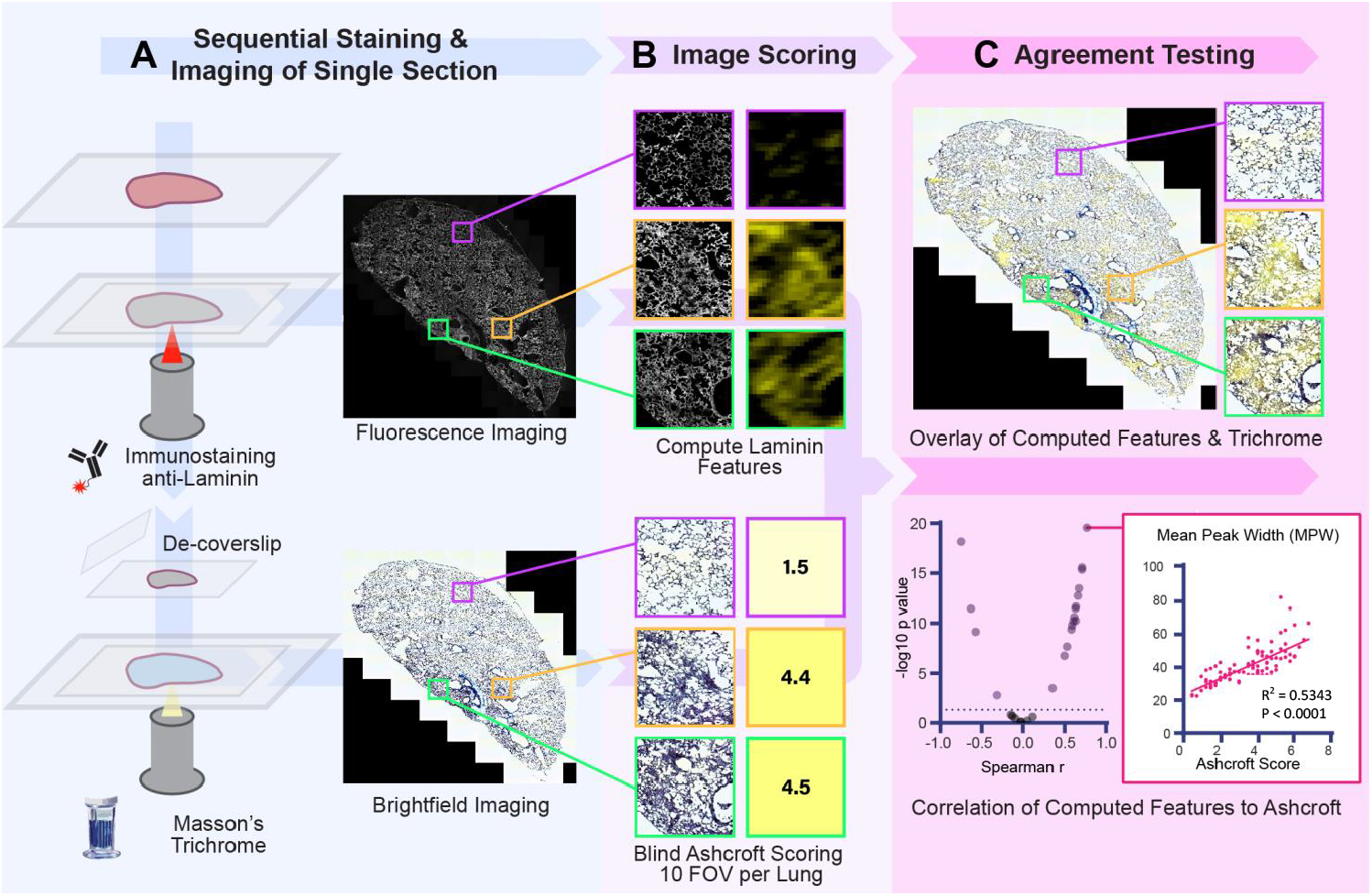
Validation of Quantitative Remodeling Scorer: Association of Computed Laminin Features to Histopathologic Ground Truth. **(A)** Diagrammatic representation of the lab workflow for sequential staining of single section for laminin and subsequently Masson’s trichrome. **(B)** Two scoring modalities performed on the immunofluorescence and brightfield acquisitions, respectively. Immunofluorescence scoring **(B, Top)** is performed on subsampled tiles of the larger image, and consists of multiple independent computational measures on per-pixel values and multi-pixel relationships within each tile. Tiles are color-coded yellow based on their score, with higher yellow values indicating higher score. Histopathologic scoring via Modified Ashcroft **(B, Bottom)** is performed on simulated 10x Fields of View (FOV). Each field is given a mean score from four independent, blinded scorers. **(C, top)** An overlay of the brightfield trichrome image with the computed laminin feature map. **(C, bottom left)** Volcano plot of the r and p values for the spearman correlation testing on the various computed laminin features against the mean Ashcroft score **(C, bottom right)** The simple linear regression of the highest scoring laminin feature chosen for the quantitative remodeling scorer, Mean Peak Width (MPW) is shown. n = 10 lungs, with 9 or 10 random fields selected per lung (98 fields total) for histopathologic scoring and subsequent comparison and regression analysis.

Scoring of images across modalities is demonstrated in Figure 2B. Image processing of laminin immunofluorescence proceeds as follows: image is subsampled into tiles, and laminin feature scores computed on each tile independently. Features scores are defined as a singly-parametrized analysis, such as gross fluorescence intensity, computed image textures including several Haralick features, and multiple line profile/histogram analyses. A chosen feature is visualized by applying a LUT to a normalized range of feature scores and generating a heatmap from those tiles. In this way, additional channels are directly appended to the original fluorescence image, with each channel representing a single feature score. Examples of these computed heatmaps can be seen in Figure 2B, top. For traditional histopathologic scoring, the Modified Ashcroft scoring system was used. Simulated single fields of view (∼900 microns square) were shown to trained and blinded scorers. The final histopathologic score for each field is the mean of the four scorers, as demonstrated in Figure 2B, bottom.

Finally, a test of association was performed between the histopathologic score and the mean laminin feature score for the same 900 micron field (airways and other non-parenchymal tiles excluded from the average). An overlay of the heatmap onto the Masson’s trichrome can be seen in Figure 2C, top. Spearman’s correlation between all computed laminin features and modified Ashcroft scores indentified in many significant features being found (Figure 2C, bottom left). The highest correlation was found with the laminin feature Mean Peak Width (MPW), with a Spearman correlation coefficient (r) of 0.768 (p=10^−19^) to the modified Ashcroft scoring (Figure 2C, bottom right). Further mentions of a quantitative remodeling scorer (QRS) in this manuscript refer to a score derived from the MPW feature. A full list of the correlation of computed features to modified Ashcroft is included in Supplemental Table 1. To test the batch-to-batch reproducibility based on potential variance in tissue sample processing, two independent animal cohorts of mice (n=6 and 4) were treated with bleomycin, with start- and endpoints falling on separate days. The agreement of MPW to modified Ashcroft score across ten randomly chosen fields per lung, averaged within each independent cohort, were not significantly different and share a best-fit line (Supplemental Figure 1).

### Example use case: association of tertiary lymphoid structures (TLS) with tissue remodeling

To demonstrate an analysis enabled by QRS’s granular profiling of tissue remodeling, we chose to investigate the association of lymphocytic aggregates, frequently termed tertiary lymphoid organs or structures (TLS) with fibroproliferative tissue in the bleomycin injury model. In a multicolor immunofluorescence assay, B cells were identified in tissue via B220 (CD45R) staining. Representative micrograph can be seen in Figure 3A. TLS, defined by densely nucleated structures of B cells >100um in length along a long axis, were observed in nine out of thirteen bleomycin treated mouse lung sections at 14 days post bleomycin. These nine sections were carried forward into subsequent analyses. A QRS map was computed for each whole lung section with a resolution of 75 square microns. A heatmap of QRS on a lung section can be seen in Figure 3B, top. A distance field to the nearest TLS was computed and can be seen in Figure 3B, bottom. To control for the area of tissue (and thus number of tiles) at increasing given radii from TLS being unevenly distributed across lungs (Supplemental Figure 2), distances were binned into 25-micron intervals, and the average QRS at each radius plotted (Figure 3C). The continuous QRS was binned into ‘Healthy’ and ‘Fibrotic’ with reference to the linear regression to modified Ashcroft score described in Figure 1 (Figure 3D). With this categorical variable, we then plotted the proportion of tissue scoring as fibrotic, calculated as the quotient of tissue meeting an arbitrary threshold for fibrotic (QRS > 42, a regressed equivalent to between 4 and 5 on the modified Ashcroft scale) to all tissue at a given radius (Figure 3E). A final analysis finds significant differences in the median distance of ‘Fibrotic’ tissue to TLS when compared against ‘Healthy’ tissue from the same section (Figure 3F).

**Figure 3.**
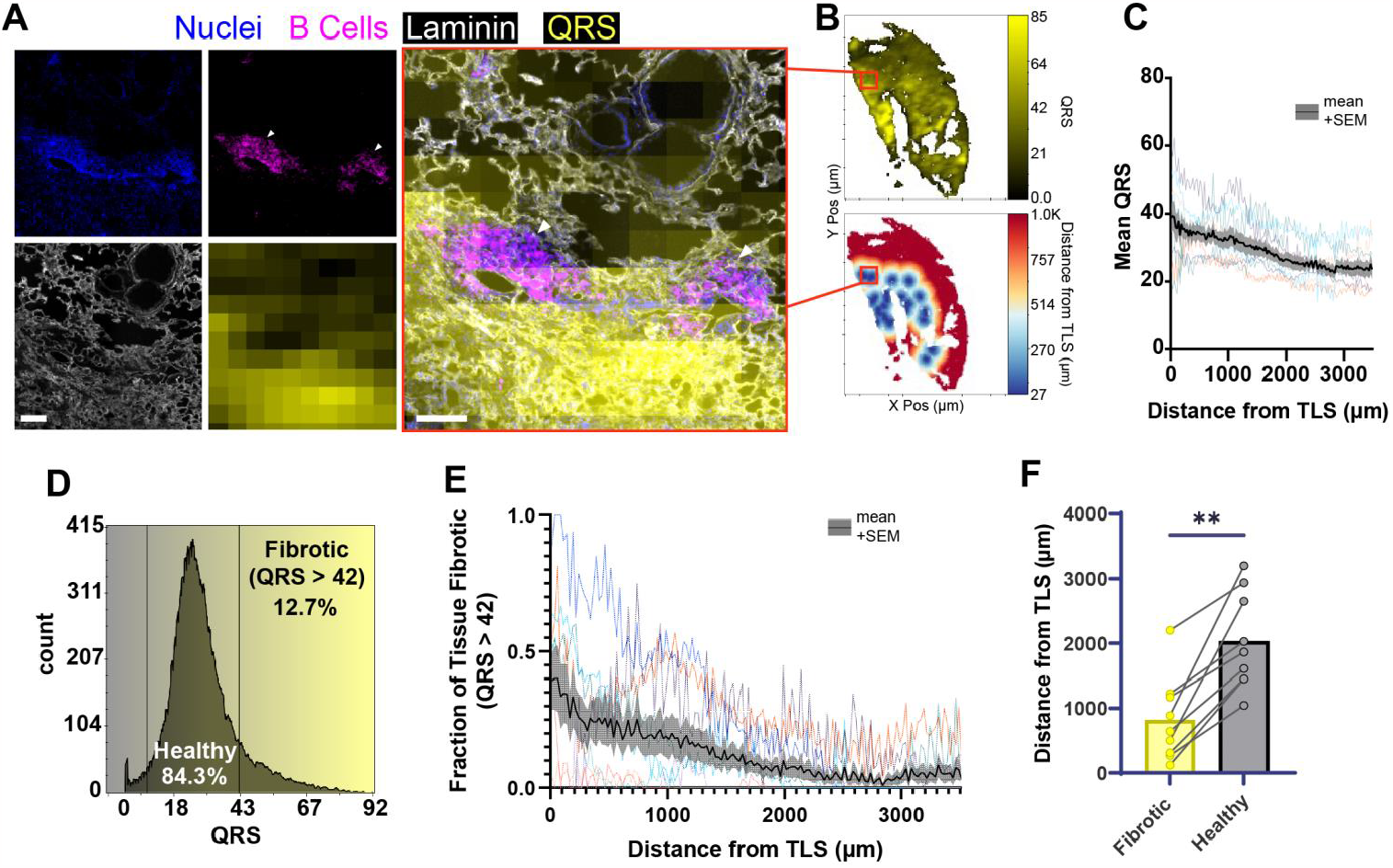
Quantitative Remodeling Scorer (QRS) Enables Testing of Spatial Hypotheses: Fibroproliferative Tissue Associate with Tertiary Lymphoid Structures (TLS). **(A)** Representative micrograph of multicolor immunofluorescence panel + QRS heatmap. Nuclei (NucGreen), B Cells (anti B220), laminin (anti-Laminin) and QRS heatmap (computationally derived from laminin) are visible as single channels **(A, left)**, and composite overlay **(A, right)**. White triangles indicate tertiary lymphoid structures. Scale bar = 100 microns. **(B)** Parametrized tiles visualized using standard cytometric software, with the field from (A) bounded in red. Heatmap of whole section QRS **(B, top)**. Distance field indicating distance from any tile to the nearest TLS **(B, bottom). (C)** Plot of the mean QRS at a given distance to the nearest TLS (in 25 micron bins). Individual colored lines represent individual lungs, black shaded line is the overall mean + SEM. **(D)** Histogram of tissue QRS, with regions gated as ‘Healthy’ and ‘Fibrotic’ at a QRS of 42. **(E)** Plot of the fraction of all tissue scoring fibrotic (QRS > 42) at a given distance from the nearest TLS. Individual colored lines represent, black shaded line is the overall mean + SEM.) **(F)** Scored lung sections split into ‘Healthy’ (QRS < 42) or ‘Fibrotic’ (QRS > 42). Plotted median distance of ‘Healthy’ or ‘Fibrotic’ tissue from nearest TLS. Scored lung sections are paired. Bars indicate the mean within each group. (n = 9, Wilcoxon matched-pairs signed rank test. **p < 0.01)

## Discussion

The study of disease is undergoing a revolution of ‘spatial’-omics. Whereas in the past, highly multiplexed data would require dissociation of tissue, the current proliferation of methods is enabling these massively parametrized assays to function *in situ*, preserving essential spatial context. However, and likely due to methodologic inertia from cytometric techniques, these methods tend to exclude any analysis of the tissue morphology and composition. In short, there is a disparity between our capacity to collect data with modern multiplex imaging and our ability to extract useful, testable data from it. This project focuses on one aspect of that disparity: the ability to quantitatively define the level of tissue remodeling across large immunofluorescent acquisitions. We compare this algorithm, named QRS, favorably against the current gold-standard for quantitatively scoring pulmonary fibrosis in experimental models.

The final feature chosen for the QRS, mean peak width, can be related to the common lung morphometric measure of chord length. While chord length measures the length of empty airspace between parenchymal tissue, mean peak width serves to measure the thickness of the interstitial basement membrane itself, and ignores empty space. By taking the mean of many line profiles drawn through a tile, an average thickness of interstitial space in that tile can be determined. A benefit of this approach is its inherent robustness to differentials in processing; poorly inflated or locally underinflated regions of lung are differentiated from genuine tissue remodeling. Similarly, peak width does not directly rely on fluorescence intensity, provided the laminin signal is sufficiently above the noise floor. Sensor overexposure to the point of signal bloom into adjacent pixels, and/or tissue outside the focal plane can result in artificially broad laminin peaks and will increase the score. Exposure management is trivial for trained microscope operators, and out-of-plane issues can be ameliorated by acquisition via confocal, structured illumination, or other optically sectioning imaging modalities. Widefield imaging, as shown in this manuscript, is fit for task provided tissue thickness is less than the depth of field for the imaging configuration. As always, consistent processing, staining, and the inclusion of fluorescence controls are required for any analysis.

Future enhancement of the tool includes a focus on optimizing computational runtime and integration into existing histopathologic tools. Certain features which scored highly in our agreement testing, such as the Haralick feature informational measure of correlation 2 (IMC2), are already implemented in software packages such as Qupath^18^, and can be computed much more quickly than the mean peak width feature used in the above analyses. It is up to the user to weigh software familiarity and interoperability with existing workflows against the strength of association between laminin features and the modified Ashcroft rubric. The concept of identifying profiles of healthy tissue structure to better identify areas of remodeling can likely be easily extended into other domains of biomedical image analysis.

Network analyses could identify where well-defined parenchyma is dysregulated by failing to identify the regular branched architecture in those regions. More complex computational methods such as computational neural networks (CNNs) may provide more granular phenotyping of tissue with the potential downside of methodologic opacity.

This tool addresses the growing need for high-throughput computational morphometric methods in lung histology. It provides reproducible, continuous quantification of tissue remodeling at a user-defined spatial scale, reducing the labor and subjectivity involved in human-scored systems. Immunofluorescence multiplexing enables the testing of hypotheses dissecting the spatial heterogeneity of biology in fibroproliferative lung disease.

## Supporting information

Manuscript Supplement

## Acknowledgements

BPC design, development, implementation and testing of algorithm and computed feature scoring; RTH brightfield and immunofluorescence imaging, statistical testing; BPC RTH NB JMS modified Ashcroft scoring; BPC RTH JMS manuscript writing, editing, and figure generation. Thomas Barker for use of computational resources. Melissa Brevard in the Cardiovascular Research Center (CVRC) histology score for assistance with histochemical stains. This work used the Leica Thunder TIRF epifluorescence microscope in the Advanced Microscopy Facility which is supported by the University of Virginia School of Medicine, Research Resource Identifiers (RRID): SCR_018736. Funding from NIH UL1TR003015 and KL2TR003016.

