## Supplementary material for "Local, Quantitative Morphometry of Fibroproliferative Lung Injury using Laminin": Manuscript Supplement

**Supplemental Material**
**Figures**

**SUPPLEMENTAL FIGURE 1**

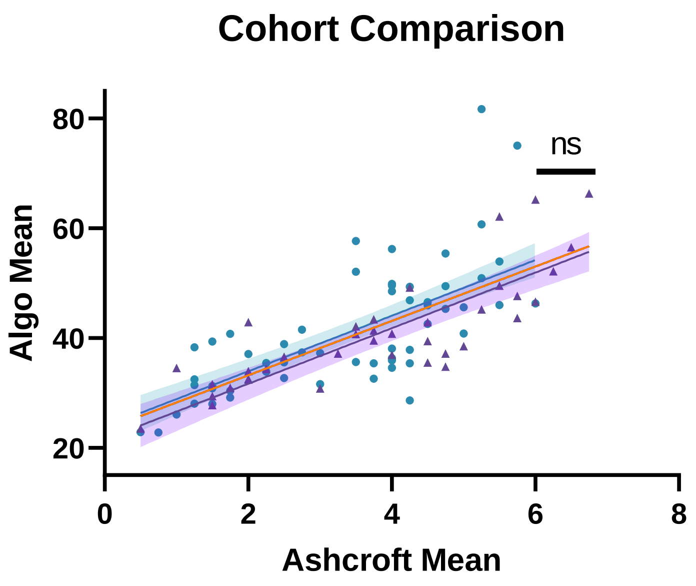


**Supplemental Figure 1. Comparison of fits across two independent experimental cohorts.** Comparison of two fit lines between the QRS and modified Ashcroft scoring using the Extra sum-of-squares F test indicates no difference in the lines.

**SUPPLEMENTAL FIGURE 2**

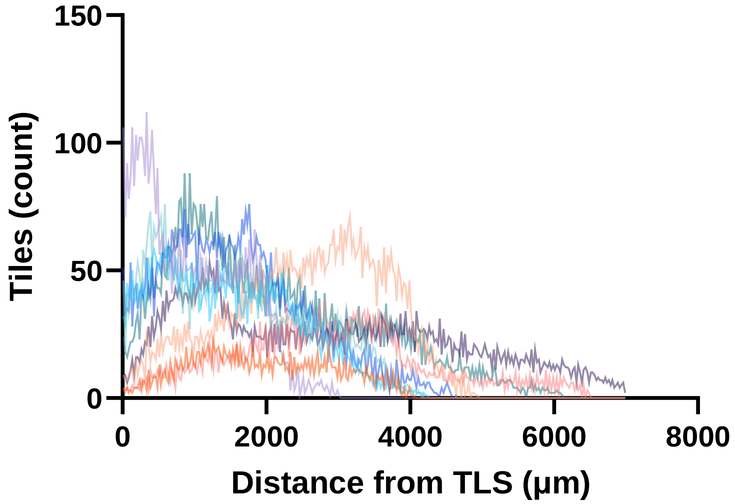


**Supplemental Figure 2. Tile Count as a Function of Distance from Tertiary Lymphoid Structures (TLS)** Histogram of tile frequency vs distance from the nearest TLS, with each color a midline sagittal section from independent mice. Differences in lung morphology and injury phenotype results in highly variable frequencies of tissue at a given distance from a TLS.

| Comparator | source of data | Analysis | Spearman r | p value | Spearman r abs |
|---|---|---|---|---|---|
| Algo Mean | laminin IF | QRS Application | 0.767941 | 2.81E-20 | 0.767941 |
| Angular Second Moment | laminin IF | QuPath Haralick Feature | -0.74933 | 7.02E-19 | 0.74933 |
| Information Measure of Correlation 2 | laminin IF | QuPath Haralick Feature | 0.709895 | 2.75E-16 | 0.709895 |
| Algo St Dev | laminin IF | QRS Application | 0.672565 | 3.35E-14 | 0.672565 |
| Sum Entropy | laminin IF | QuPath Haralick Feature | 0.63501 | 2.18E-12 | 0.63501 |
| T150 Mean | laminin IF | QRS Application with higher intensity threshold | 0.633857 | 5.72E-11 | 0.633857 |
| Entropy | laminin IF | QuPath Haralick Feature | 0.630418 | 3.5E-12 | 0.630418 |
| Difference Variance | laminin IF | QuPath Haralick Feature | -0.63013 | 3.6E-12 | 0.63013 |
| T160 Median | laminin IF | QRS Application with higher intensity threshold | 0.615353 | 2.54E-11 | 0.615353 |
| T160 Mean | laminin IF | QRS Application with higher intensity threshold | 0.605123 | 6.57E-11 | 0.605123 |
| Lam Mean | laminin IF | Laminin fluorescence average | 0.58135 | 4.27E-10 | 0.58135 |
| Inverse Difference Moment | laminin IF | QuPath Haralick Feature | -0.5725 | 7.32E-10 | 0.5725 |
| Correlation | laminin IF | QuPath Haralick Feature | 0.528783 | 2.18E-08 | 0.528783 |
| Sum Average | laminin IF | QuPath Haralick Feature | 0.497984 | 1.81E-07 | 0.497984 |
| Difference Entropy | laminin IF | QuPath Haralick Feature | 0.355582 | 0.000327 | 0.355582 |
| Information Measure of Correlation | laminin IF | QuPath Haralick Feature | -0.31403 | 0.001639 | 0.31403 |
| Lam Std Dev | laminin IF | Laminin fluorescence | -0.14529 | 0.155626 | 0.14529 |
| Contrast | laminin IF | QuPath Haralick Feature | -0.12783 | 0.209709 | 0.12783 |
| Sum of Squares | laminin IF | QuPath Haralick Feature | 0.116628 | 0.252766 | 0.116628 |
| Lam Min | laminin IF | Laminin fluorescence | -0.09544 | 0.352405 | 0.09544 |
| Sum Variance | laminin IF | QuPath Haralick Feature | 0.049632 | 0.627442 | 0.049632 |
| Lam Max | laminin IF | Laminin fluorescence maximum | -0.0271 | 0.792166 | 0.0271 |

### SUPPLEMENTAL TABLE 1

**List of Computed Laminin Features for Spearman Correlation against Modified Ashcroft**

List of laminin immunofluorescence-derived comparators in the spearman correlation, sorted by absolute Spearman r.

### QRS APPLICATION

The tool exists as a standalone Python application, where a user can search for compatible images through a graphical user interface (GUI) and run the scoring algorithm. Only CZI or TIFF images are currently supported. The user must input several important parameters in the GUI including tile size, the number of line profiles, the laminin channel number for multi-channeled images, a pixel intensity threshold, and an output file path. Using these parameters, the program first selects the laminin channel and subdivides the whole slide image (WSI) into tiles of desired size. Within each of these tiles, it draws a user-defined number of evenly spaced line profiles in both the X and Y dimensions. These line profiles are filtered with SciPy filt-filt and the smoothed plot is used to capture the width of the laminin network, which we observe to correlate highly with the progression of fibrosis as determined by modified Ashcroft score. At a user-defined or algorithmically identified threshold, we calculate the mean width under the peaks in the line profile, which is referred to as Mean Peak Width (MPW). The program classifies a point where the line profile crosses threshold as either crossing up or crossing down. It then correctly pairs the crossing points with either each other, the beginning, or the end of the tile depending on the sequence of up/down crossings. This ensures that it only calculates the width under the line profile (i.e. the width of true signal, not the width of background signal). The scores for each line profile are averaged into a score for the entire tile. There is an optional feature to then smooth the tile-map by taking an average with neighboring tiles in all directions. The tile-map is then saved as an additional channel on a copy of the input image, which can be viewed as a composite heatmap over other image channels. From here, various downstream analyses can take place (Figure 1). The tool is capable of batch processing large sets of images, exists as a standalone application (meaning all dependencies are included in the application), and transfers all image metadata between input and output images.

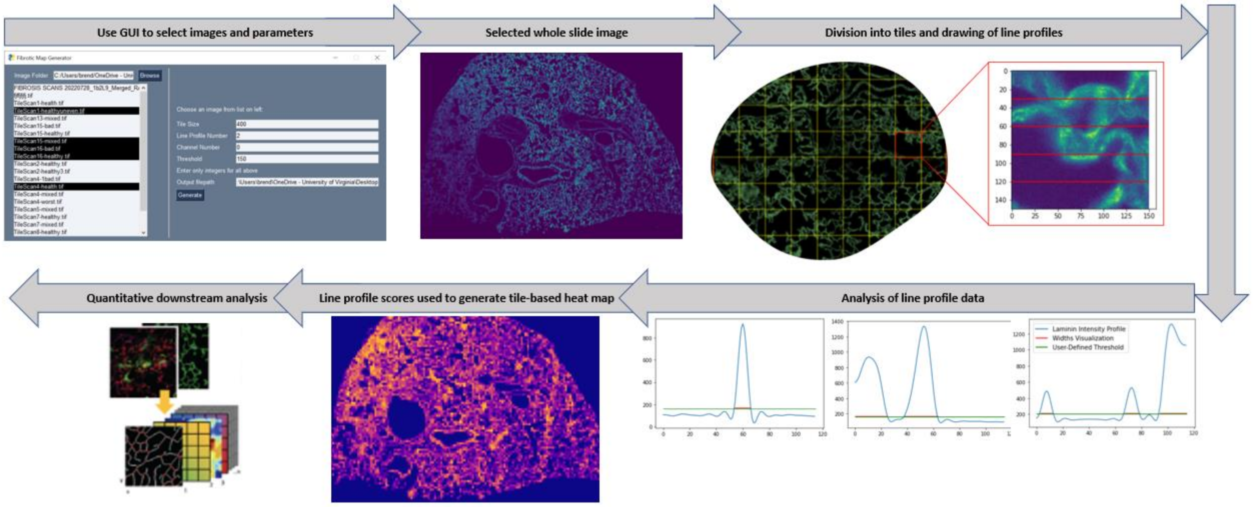


***Supplemental Figure 3. Computational process outline. Images are selected and user-defined elements are set. The tool then divides the image into tiles and generates a heat map based on the MPW score for line profiles within that tile.***

### <u>Using the Tool</u>

The tool exists as a standalone application. It can be downloaded and used without any external software, although some form of imaging software such as QuPath or ImageJ will be required to view the output image. Users will interact with the GUI to run the program and must understand what parameters to enter. Upon running the application, the user will be prompted with this screen:

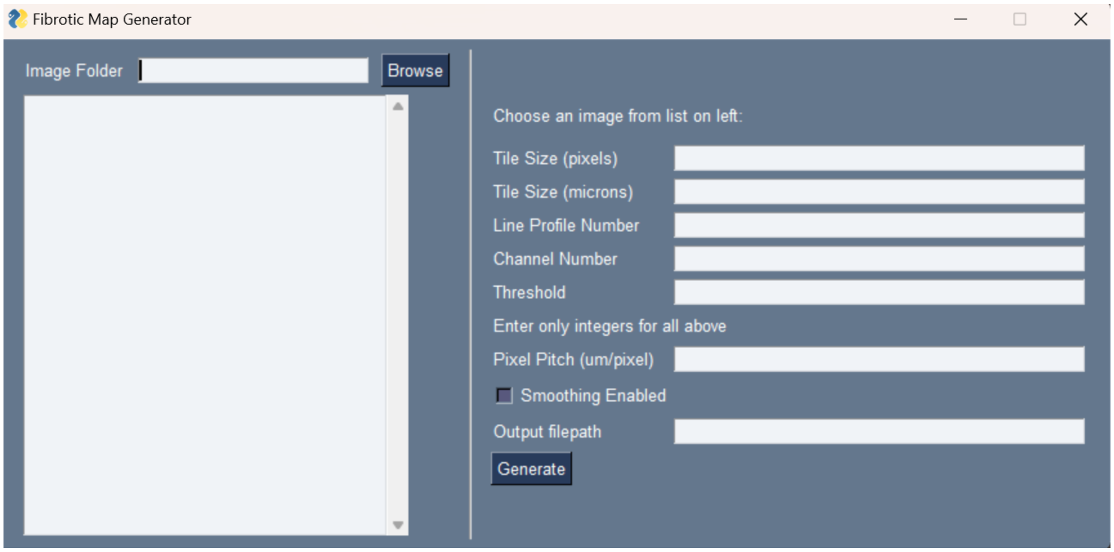


First, click “Browse” and the program will open a file browser. Select the folder containing your images of interest. Note that these images may not appear in the file browser but will appear in the list in the GUI. When you have correctly selected the image folder, the listbox (left) will populate with any compatible images stored there. Only CZI of TIFF image types will appear in the listbox. Users can also manually copy a complete file-path into the Image Folder textbox.

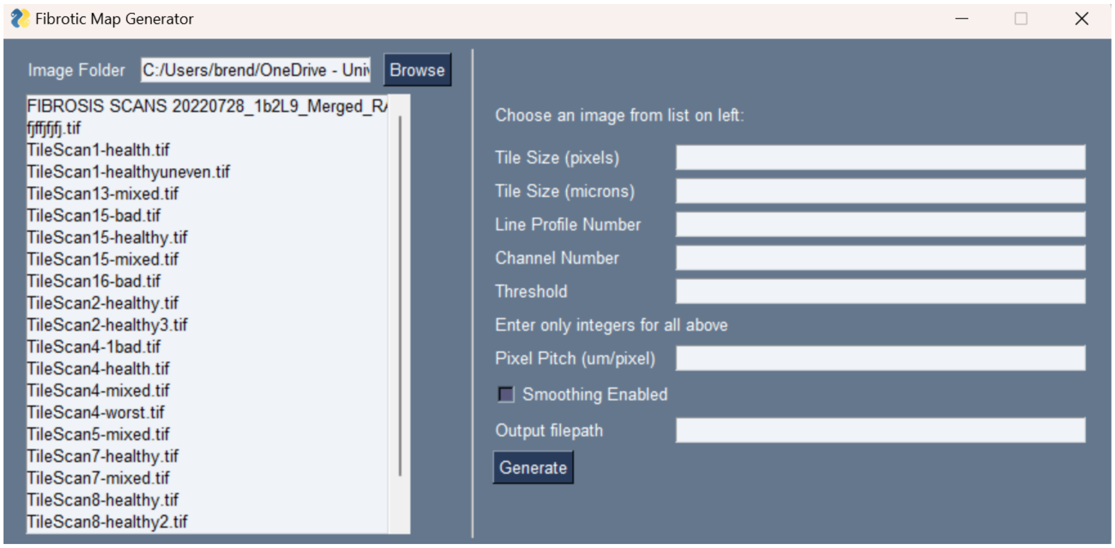


The next step will be to specify the size of the tiles. Users can either enter a pixel value or a micron value and pixel pitch. If using microns, the program will not work unless you also specify pixel pitch. Users are allowed to enter both pixels and microns, but if the values do not match the program will prompt…

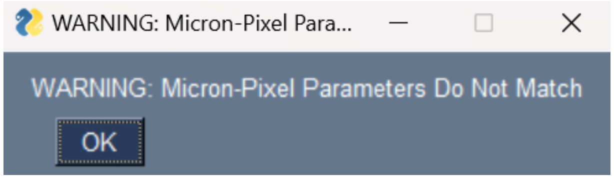


If the user closes this warning, the program will default to the given pixel value (not micron) and proceed. The user will have to end the program and reenter values to change them.

Next, the user will have to specify the number of line profiles to be drawn for each tile. Do not specify a number of profiles larger than the tile size (in pixels). The program will evenly space the number of line profiles requested across the tile. If the number of profiles is equal to the tile size, line profiles will be drawn at every pixel in the image. This is not recommended. Generally, one profile for every ten pixels is sufficient.

The channel number is used to direct the program to the laminin channel for multi-channeled images. Note that even if the image has only one channel, the user still must specify “0” for channel number. It is also important to remember that the channel number is indexed from 0. So, a three channeled image would have channels 0, 1, and 2.

The most vital input is the threshold. Shown below as the green line, this can be understood as the user’s definition of where true signal begins. If implementing background subtraction or otherwise normalizing pixel values, this parameter must still be specified as “0”. Otherwise, it is recommended that the user use a histogram of pixel intensities across their image to select the threshold. The threshold will be the value at which the program measures the width, as shown below.

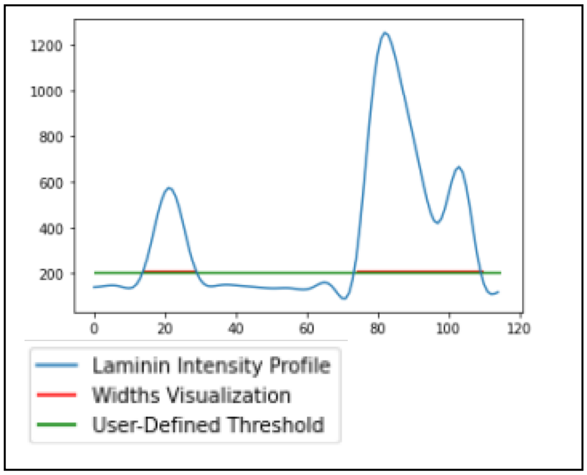


The next input, pixel pitch, only has to be entered if the user is specifying tile size in microns. It can otherwise be left blank. Please use decimal notation (ex. 0.321 um/px) The smoothing checkbox is an optional feature that makes each tile’s score an average of the scores for all adjacent tiles. It creates a more appealing heatmap and can further mitigate random sources of error from anomalous tiles at the cost of spatial resolution. Finally, the user must specify an output directory. This is typically a local folder. Please enter the entire file path, not just the name of the folder. There is not a file browser associated with the output directory. Copy and paste the file path from File Explorer for your desired output location. The program saves a new copy of the entire image, including all channels, with an additional channel being the scored heatmap. It is therefore important to check that there is sufficient storage on the device, especially for large batch processing.
